# Nanopore sequencing enables multigenic family reconstruction despite highly frequent PCR-induced recombination

**DOI:** 10.1101/2022.11.14.516392

**Authors:** Alice Namias, Kristoffer Sahlin, Patrick Makoundou, Iago Bonnici, Mathieu Sicard, Khalid Belkhir, Mylène Weill

## Abstract

This study developed a new bioinformatics pipeline to acquire all the different copies of multi-copy gene families based on Oxford Nanopore Technologies sequencing of PCR products. We used this pipeline to acquire the sequences of highly similar copies of the *cidA* and *cidB* genes present in the genomes of *Wolbachia pipientis* (wPip) bacteria infecting the cells of *Culex pipiens* mosquitoes. The approach is based on read mapping, SNP calling and haplotyping, using our already wide existing reference database for the *cid* genes obtained by cloning and Sanger sequencing. We addressed problems commonly faced when using mapping approaches for multi-copy gene families with highly similar variants (or haplotypes). In addition, we confirmed that PCR amplification causes frequent chimeras which have to be carefully considered when working on families of recombinant genes. We tested the robustness of the pipeline through a combination of analyses of simulated reads and of gene sequence acquisitions through cloning and Sanger sequencing. For genes of which the haplotype cannot be reconstructed from short reads sequencing, this pipeline confers a high throughput acquisition, gives reliable results as well as insights of the relative copy numbers of the different variants.

## 1. Introduction

Although the role of gene duplication-amplifications (GDA) is now well recognized as a mechanism of adaptation to environmental variations [1], studying the structure of GDA remains a challenge. Indeed, GDA do not create any sequence novelty, with the exception of the sequence of the insertion breakpoint, and finding such restricted features is almost impossible when working with short reads sequencing. In recent years, methods such as whole genome DNA arrays and high-throughput sequencing, increased the general detection of amplifications [2]. Duplications can be identified through coverage variations in next-generation sequencing (NGS, e.g. [3]) or through real-time quantification PCR (qPCR) when targeting a specific locus. While some duplications are homologous (i.e. all duplicated copies are identical), diversification can occur following GDAs, leading to the presence of multiple polymorphic copies in a given genome. In that case, a further challenge is to identify the variants (or haplotypes) of the different gene copies. If variations are interspersed and separated by a number of base pairs larger than the sequencing read length, haplotype reconstruction using short reads is complex, if not impossible. The recent expansion of long read sequencing is thus of key interest to sequence such gene families. Yet, if long reads sequencing methods have been widely used for analyses of genomic structural variations, their high error rates (6-8% for MinION sequencing, [4]) and/or financial cost (for Pacific Biosciences sequencing) have been a major obstacle to their use for accurate variant identification. Lately, PacBio long reads have successfully enabled to identify new isoforms from RNAseq data, thanks to new tools [5]; improvement of the Nanopore basecallers with either Nanopolish [6] or Nanocaller [7] made possible the use of Nanopore sequencing for identification of inter-individual single nucleotide polymorphisms (SNPs) and variations in gene copy numbers [8], and error correction enabled using Nanopore technology for reference-free transcriptome analysis [9]. Nanopore sequencing has also recently been used for multiplex amplicon sequencing, decreasing the financial cost by 200x as compared to Sanger sequencing [10]. A persistent problem for identification of new genetic variants is the non-random distribution of long read sequencing errors, these errors being more frequent in homopolymer regions (representing approximately half of sequencing errors) and in GC-rich regions [4], making it hard to discriminate true mutations from sequencing errors even when increasing the depth of sequencing coverage. This can be mitigated by a previous knowledge of the within-gene polymorphism distribution, e.g. with a good pre-existing database. A further issue arises after having identified SNPs, when haplotype phasing is required to obtain the haplotypes. Indeed, common haplotype phasing tools (e.g. GATK, WhatsHap) require a previous knowledge on the expected number of different gene copies (copy number for multi-copy gene families, ploidy in most of the cases). This is an issue in the case of multi-copy gene families, since many of them display among-individual copy number variations, making the prior specification of copy numbers impossible.

Here, we developed a method to sequence gene families based on Nanopore sequencing of PCR products for which a database of existing variants is available. This was done in order to work on two cytoplasmic incompatibility factors, the *cidA* and *cidB* genes (altogether named *cid*), present in the endosymbiotic bacteria *Wolbachia pipientis* (wPip) genome in *Culex pipiens* mosquitoes. These genes encode key proteins involved in cytoplasmic incompatibility (CI) [11,12], a well-studied conditional sterility induced by *Wolbachia* in its arthropod hosts which is currently envisaged or implemented as a means of mosquito vectors and agricultural pests control [13–16]. Both genes, described as a tandem, are amplified and diversified in wPip with up to 6 different copies described within a single *Wolbachia* genome, the set of *cidA/cidB* gene copies present in an individual being called a repertoire [17]. Copy numbers vary among strains of wPip. The different copies of *cid* genes (also named variants) have been shown to correlate with incompatibility patterns [17,18]. Being able to obtain individual’s *cid* repertoires is thus a prerequisite to understand and predict CI patterns and evolution in mosquito populations. More than 30 variants of *cidA* and *cidB* have been previously described in wPip infecting *Culex pipiens* mosquitoes from all around the world [17–20]: the variants present in different *Wolbachia* strains were amplified by PCR using generic *cidA* and *cidB* primer pairs. The PCR products were then cloned and bacterial clones were sequenced by Sanger sequencing. This method is extremely time consuming to set up and thus inappropriate for large scale studies. Furthermore, it has limited descriptive power: due to time and money, a maximum of 48 clones were sequenced per individual PCR, making it likely to miss a rare variant in *Wolbachia* having numerous and unequally distributed variants. Despite this short coming, analysis of the variants led to observe polymorphism hotspots, and to restrict variation analyses to two specific gene regions [17–20].

Based on this existing reference database, we developed a bioinformatics pipeline to acquire repertoires using Nanopore sequencing of PCR products. Overall, this pipeline assigns each read to its closest known reference through mapping, then uses a combination of SNP calling and haplotype phasing to identify potential new variants. We validated and fine-tuned the pipeline and its efficiency to accurately find new variants in spite of Nanopore sequencing errors using a combination of read simulations and molecular biology approaches. Thresholds to discriminate true new variants from sequencing and/or PCR errors were established through PCRs targeting specific regions in order to test for their actual presence/absence. Read simulations were used to (i) confirm that the pipeline could recover reads in spite of Nanopore sequencing errors and (ii) ensure that it was possible to identify distant new variants which were absent in the existing database. The repertoires obtained were further verified through variant specific PCRs, along with cloning and Sanger sequencing, highlighting highly frequent PCR-induced recombination, causing numerous artifactual variants in both Sanger and Nanopore sequencings. Protocol changes were implemented to mitigate these recombinations. Moreover, we showed that such sequencing could give access to the relative copy numbers of the different variants.

## 2. Results

### 2.1. Pipeline overview

Our goal was to create a bioinformatics pipeline to reconstruct multi-copy gene families with variable copy numbers through Nanopore sequencing. To this aim, we worked on the *cidA* and *cidB* genes of wPip *Wolbachia* infecting various *Culex pipiens* isofemales lines.

The pipeline is based on long reads obtained from Nanopore sequencing of PCR products of *cidA* and *cidB* genes. Different PCR products can be pooled for sequencing and separated for downstream analyses (here, we pooled *cidA* and *cidB* PCR products). Since we knew that *cid* genes were made of two variable parts (one upstream and one downstream), we created artificial references by combining known upstream and downstream regions to complete the existing database (Fig 1). The reads are then mapped on the full set of references (true and artificial) to identify the references covered above a pre-determined threshold (detailed below). Mapping is done using minimap2 [21] at all steps. References above the coverage threshold are then further filtered by removing highly similar references. We remove references having less than a user-defined number X of SNPs to any other above-threshold reference, thus solving downstream SNP calling issues leading to low coverage and spurious indels (X threshold is a pipeline parameter, default 3, samtools 1.15 is used in the pipeline, [22]). Then, reads that mapped to at least one of the references in the pre-existing database are re-mapped to our selected reference subset, and SNP calling is performed using samtools mpileup (samtools 1.15). Finally, if SNPs are called, gene variants are identified through either consensus assembly or haplotype calling using either samtools consensus [22] or WhatsHap [23] (Fig 1).

**Figure 1.**
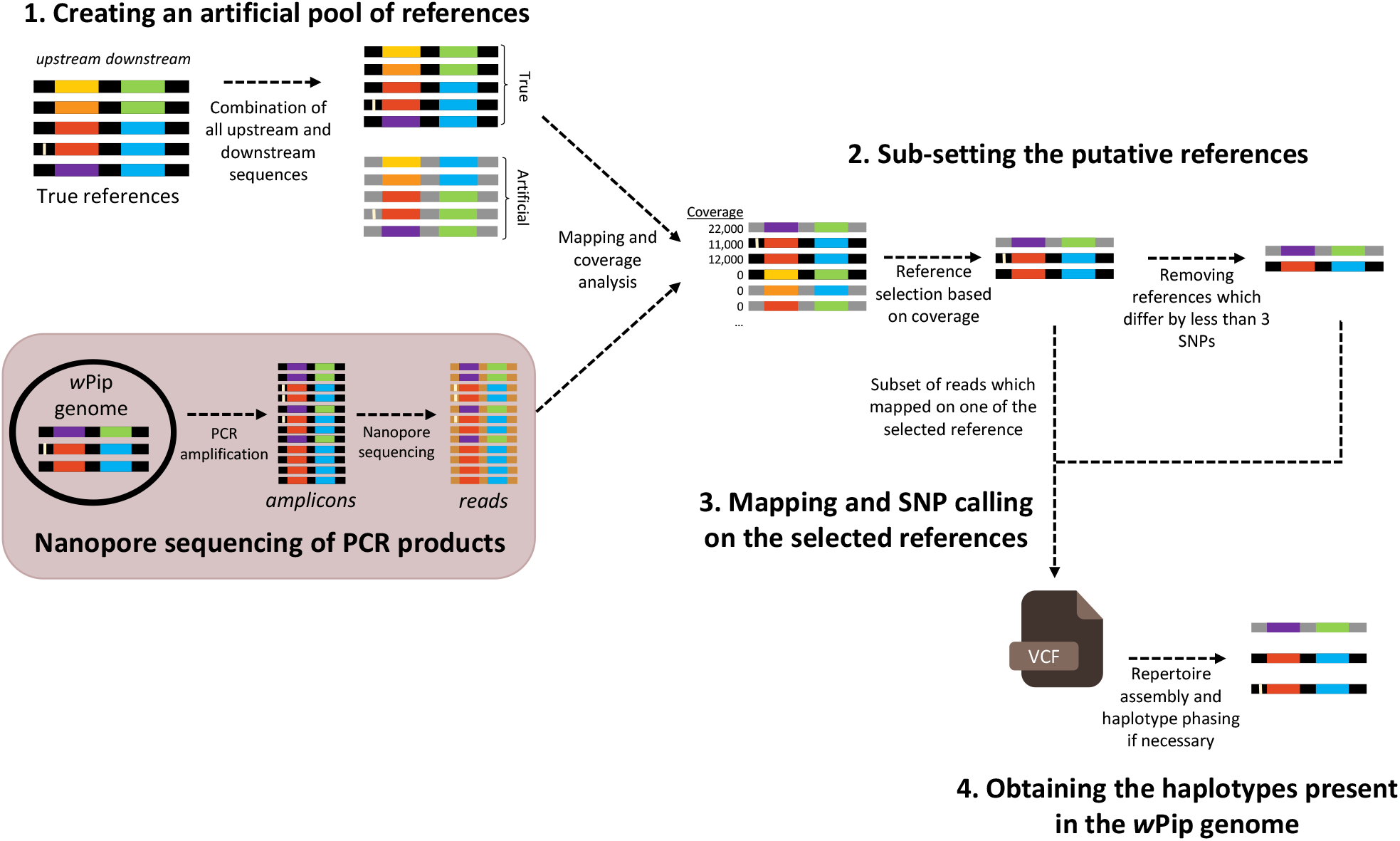
Overview of the bioinformatics pipeline. *cid* gene variants are represented with their upstream and downstream variable regions shown by different colors. A black background represents the true references, a grey one represents artificially created variants, and a brown one Nanopore sequencing reads. Some variants differ by few SNPs only, outside of colored variable regions. SNPs are represented by a clear bar.

While the pipeline requires a pre-existing database, new variants can be recovered through SNP calling.

### 2.2. Definition of the threshold for true variants by specific PCRs

One of the key steps was to determine if all identified variants were truly present in the genome of the studied wPip strain or if some were artifactual. For all the wPip strains analyzed, the coverage distribution across all references showed that most references had no coverage, few had a high (more than 10% of reads) coverage, and some had a low-to-intermediate coverage (see an example in Fig S1). In order to determine whether those “low-to-intermediate coverage” variants were truly present in the sample, we designed multiple specific primers to test the presence of these variants by PCR, in the exact same DNA matrix used for Nanopore sequencing. We only investigated cases for which specific primers of the questioned variant could be designed (i.e. primers amplifying solely the target variant and no other variants present in the repertoire, Fig 2). We further restricted tests to cases for which both a negative control (i.e. an infected *Culex* mosquito for which the primers should not amplify any variant), and a positive control (i.e. an infected mosquito for which the primers should amplify a variant, Fig. 2) could be used. We considered that a variant was truly present in a strain, and that the strain could thus be used as a control, when the variant was highly covered in Nanopore sequencing and previously found by Sanger sequencing [17–20]. This design enabled to eliminate the potential non-specificity of primers, a problem that can be frequent on *cid* variants due to the recombinant nature of the genes.

**Figure 2.**
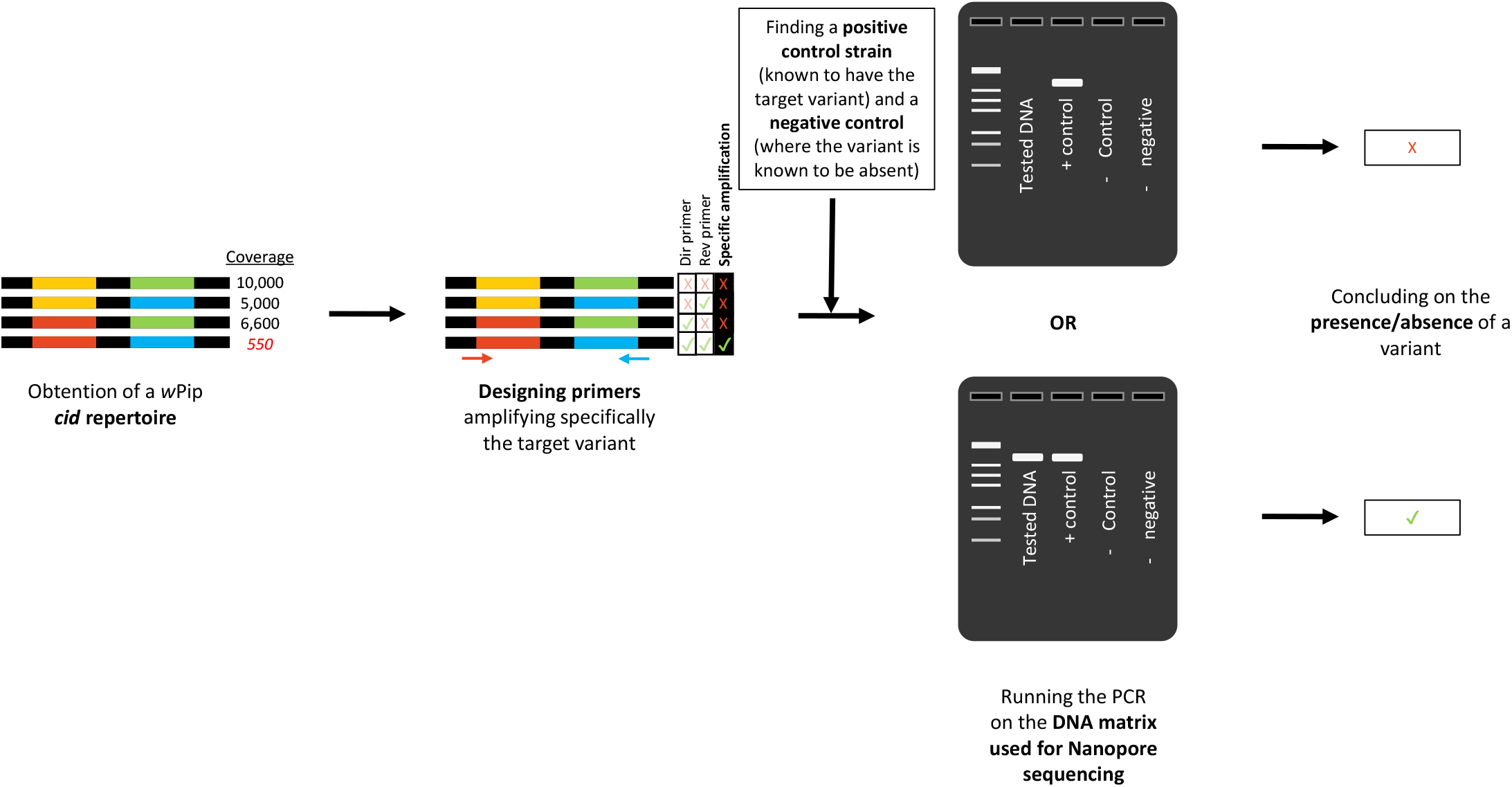
Description of the process to test for the presence/absence of a specific variant using specific PCRs. Primers amplifying this specific variant and not others have to be found, then true positive and negative controls are used (other strains in which this specific variant is specifically present or absent).

Overall, we successfully tested the presence/absence of 8 “low to intermediate-coverage” variants, in 8 different wPip strains repertoires and found that none of them could be amplified by PCR (shown in Table S1, either in dark blue or in orange, depending on whether the variant had been found in Sanger or not). Since the number of reads differed among wells, we established a relative coverage threshold, expressed as a percentage of the total number of reads. Using specific PCRs, we found that truly present references were those covered by at least 10% of the total number of reads, or a depth above 1,500 reads.

### 2.3. Subsetting the putative references to prevent SNP calling issues

Here, since we are using targeted long reads, the reference and the reads have the same length. We thus expect the average coverage to reflect the depth at each position. We used samtools 1.15 [22] for SNP calling, and found that keeping references which are really close (differing by a few SNPs) caused SNP calling problems, by artificially introducing INDELs and diminishing the coverage. The conclusion that SNP calling issues are involved, rather than mapping issues was reached because using samtools depth and samtools mpileup (also giving the depth at each position for each reference, along with calling SNPs) on the same .bam file and with the same options (setting the minimum quality at 13 for both, and increasing the maximum depth value to 70,000 for mpileup) gave drastically different results: the coverage obtained with samtools mpileup can be below 100 reads, when samtools coverage outputted a coverage of several hundred reads). This issue was solved by keeping a single representative of each pair of highly similar references.

Above-threshold references were thus subsetted to keep a single representative of each pair of references differing by less than three SNPs. We found that a 3-SNP threshold was sufficient to get rid of SNP calling issues.

### 2.4. Simulations confirm pipeline’s ability to recover variants

Although Nanopore sequencing quality has drastically improved in the last years, the rate of sequencing errors is still estimated to be around 7% [4]. We tested, using simulated reads, the ability of the pipeline to recover all variants present in spite of Nanopore errors. To our knowledge, Nanopore read simulators such as NanoSim [24], DeepSimulator [25], SimLord [26], and SNaReSim [27] are designed for genomic data and do not mimic targeted sequencing where the majority of reads covers the whole amplicon. We therefore wrote our own read simulator for targeted data based on the full-length transcriptome read simulation pipeline in [9] (isONcorrect) (details of the simulation script in the method section Nanopore read simulation). Using our simulation script, we simulated 20,000 Nanopore reads from existing references. We tested the influence of (i) the read error rate, (ii) the number of distinct variants and (iii) the repertoire’s complexity on the pipeline’s ability to recover the references. To do so, we simulated reads from variants in eight different settings. The eight settings were formed from varying error rates (4 or 12%), gene copy numbers (2 or 6 distinct copies) and repertoire complexity (starting from existing variants which are either highly similar or strongly different). The pipeline was run on reads simulated from all these cases, and the right repertoire was recovered each time. This simulation confirmed that our pipeline was robust at error rates higher than the typical Nanopore error rate of about 7%.

### 2.5. SNP calling and haplotype calling enable to properly recover new variants

We also tested the ability of the pipeline to recover true new variants (i.e. not yet present in the complete database). To that extent, we used our Nanopore read simulator, which enables to create simulated mutants with a user-set mutation rate. We created new variants by introducing a fixed number of SNPs in the existing ones. Nanopore reads were then simulated from (i) simulated variants only and (ii) a mixture of simulated and existing variants. For this experiment, reads were simulated with a 7% error rate, as we showed above that the pipeline correctly recovered the references with error rates up to 12%, and because 7% is close to recent estimates of the realistic Nanopore sequencing error rates [4]. Simulated SNPs were introduced in the existing variants in such a way that the new simulated variants differed from the existing ones by a distance equal to the median pairwise raw (Hamming) distance among true references (the database being already wide, finding a new variant differing from all others by more than this distance was highly unlikely). No INDELs were introduced since previous work showed they were unlikely in *cid* genes [17,18,20], even if the read simulation script used enables to introduce INDELs. Simulating sequences enabled us to know exactly where the SNPs were located and thus to check our pipeline’s performance in a controlled scenario.

Using 100 different simulations, we computed the true number of SNPs (ranging from 41 to 71), along with the number of SNPs recovered by SNP calling. We found that on average, 3.4 ± 3.1 SNPs were missed (mean ± standard deviation, ranging from 0 to 18). There were two distinct cases for missed SNPs: (i) SNPs located in repeated regions (e.g. AATTAATA -> AATTTAAT, likely considered by the caller as a spurious SNP resulting from a sequencing error); (ii) SNPs not called in spite of a high coverage of the alternative base (e.g. at some positions, 6500 reads supporting the alternative base, yet the SNP was not called). This last issue was not solved by increasing the prior on the substitution rate. We thus wrote an awk script to detect such issues of missed SNPs (in spite of a high number of supported reads). Using this script on our true sequencing data, we found no occurrence of such issue, suggesting it might only occur when SNPs are numerous.

### 2.6. Comparison with Sanger sequencing results

Repertoires obtained through Nanopore sequencing were compared with repertoires previously obtained by Sanger sequencing [17–20]. We found an overall agreement between Nanopore-obtained repertoires and Sanger-obtained repertoires (Table S1). A total of 54 variants were detected for *cidA* (excluding variants which were only found in low coverage in Nanopore sequencing, highlighted in dark blue in Table S1). Among those, only 4 differed between Sanger and Nanopore sequencing (excluding variants shown in dark blue, i.e. counting pink, light blue and orange cells). For *cidB*, 41 variants were detected in total, and 8 disagreements among sequencing methods were found. Out of the 12 total disagreements, 4 variants were only found in Nanopore sequencing (highlighted in pink), and 8 were only found in Sanger sequencing (or found with a low coverage in Nanopore sequencing, which we determined to be spurious, highlighted in light blue or orange). While it was expected to find some additional variants in Nanopore sequencing, since this method gives a coverage of up to 40,000 reads per gene, as compared to the analysis of 24 to 48 clones used in Sanger sequencing, the opposite case was not expected.

In order to solve these discrepancies, we used the same approach as for threshold establishment: we identified a target region which could be diagnostic of the presence/absence of a given *cidA* variant, then designed specific PCR primers and checked for its presence/absence (Fig.2). This was done on a total of 9 variants whose presence/absence differed between Sanger and Nanopore sequencing. All variants which were found only in Nanopore with a high coverage were detected through specific PCR. Conversely, none of the variants only found by Sanger sequencing were found (Table S1).

### 2.7. Artifactual variants are due to PCR-induced recombinations

Close examination of artifactual variants’ sequences found both by Sanger and Nanopore sequencings, revealed that they were all putative recombinant variants that could result from recombination of variants present in the repertoire of the same wPip strain. It seemed that more artifactual variants were observed when the wPip strain had a high number of *cid* variants (for example Slab, [19]). Sanger and Nanopore sequencings were both run on PCR products, and PCR amplification has previously been described as a potential cause of artificial recombinations, especially when using a high number of PCR cycles [28–30]. To test the putative link between artifactual variants and number of cycles used for PCR, we chose the *Wolbachia* from the Slab isofemale line because it shows many *cid* variants [19]. We amplified the *cidA* genes in 4 individuals using either 31 or 35 PCR cycles (on the same DNA for both PCRs), then sequenced the amplicons through the same Nanopore sequencing protocol. Slab had also been previously Sanger sequenced using 35 PCR cycles. We found more artifactual, chimeric reads when 35 cycles were used compared to 31, coverage of the corresponding references being up to almost 10% at 35 cycles. The *cidA* repertoire has 4 different variants, and two additional artificially created variants were repeatedly sequenced, by both Sanger and Nanopore sequencing (namely *cidA-III-gamma(3)-25* and *cidA-III-beta(2)-12*, which can come from recombinations between existing variants). Similarly, two additional artifactual *cidB* variants had been found in Sanger sequencing in Slab (using more PCR cycles), suggesting this strain may be prone to recombinations possibly due to its high number of *cid* variants.

### 2.8. Nanopore sequencing coverage predicts relative variant copy numbers

Nanopore sequencing of different wPip strains gave consistent differences in coverage between the variants (some variants being more covered than others). To test whether these differences in coverage may reflect variations in copy number among the different variants, we analyzed the three *cidA* variants (*cidA-II-alpha-15, cidA-II-alpha-7*, and *cidA-II-beta-15)* of the *Wolbachia* from the Lavar (Lv) isofemale line, which were identified both by Sanger and Nanopore sequencings. Since specific amplification of the different variants requires a sequence of approximately 600 bp, which is too long for classic quantitative PCR (qPCR), we used digital droplet (ddPCR) to amplify specifically the variants. Since different PCR primers were used for each variant, we first investigated whether these PCRs had similar efficiencies, to ensure that differences in variant quantifications resulted from true differences in copy number rather than distinct PCR efficiencies. We designed specific primer pairs for each of these variants, cloned them and validated the specificity of each primer pairs on the clones (a given primer pair must amplify the corresponding clone only to be validated). To ensure that differences in PCR efficiency did not affect differences in nanopore coverages, we ran the three specific PCRs on the same amount of DNA from each clone. We obtained similar concentrations, showing differences in Nanopore coverage indeed reflected concentration variations (Table S2A).

Since quantifying the different copies in wPip-Lv by ddPCRs required the use of total DNA containing a mixture of all *cidA* present, we had to ensure that we were able to quantify all *cidA* copies. To do so, we used three generalist *cidA* primer pairs located in a monomorphic region of the *cidA* gene that should amplify all gene variants (Table S3). We tested each generalist pair of primers using an equimolar mixture of the isolated clones. Surprisingly, and although ddPCR is supposed to be devoid of efficiency effects associated with fragment size, we found that concentration (number of copies) obtained were widely different when the fragment size was 150, 580 or 1300 basepairs (bp) (715.8, 535.4 and 307.7 copies/μL, respectively), showing that the fragment size strongly influences the obtained concentration (Table S2B). For further experiments, we chose the 580 bp generalist pair of primers as it was the pair giving (i) amplified fragments with a size close to the specific fragments (~600 bp), and (ii) a concentration close to the sum of individual copy numbers (i.e. 586 copies/μL) in the equimolar mix of the three specific clones.

We then quantified the different variants in the DNA obtained from a single Lv mosquito. Comparing the relative quantifications of the different variants using ddPCR with relative coverage obtained in Nanopore sequencing, we found an overall agreement with fewer copies of the *cidA-II-gamma-15* variant compared with the two others, suggesting that Nanopore sequencing coverage could give proper insights in relative copy numbers (Table S4). Combined with a qPCR giving the total number of *cidA* copies, such relative coverages could be used to obtain variants’ copy numbers.

## 3. Discussion

Within a genome, there can be multiple highly similar copies of a multi-copy gene family. For *cid* genes, our focal example here, we called these different copies ‘variants’, and the full set of copies per genome is called a ‘repertoire’. Sequencing and assembling such families, with highly similar copies and copy number variations is still an issue. Indeed, long reads are highly error-prone, making it difficult to differentiate true polymorphism from sequencing errors, and short reads fail to enable haplotype reconstruction. Here, we developed a bioinformatics pipeline enabling to reconstruct the full repertoire of a gene family within a single genome through long read Nanopore sequencing of PCR products, based on an existing reference database of Sanger sequences. Using numerous specific PCRs, we established the correct coverage threshold to discriminate true variants from artifactual ones. We validated the pipeline using both simulated reads and molecular biology approaches, with cloning and Sanger sequencing along with other specific PCRs, confirming its ability to recover gene repertoires in spite of Nanopore sequencing errors, both by correctly identifying previously sequenced variants and by successfully assembling new haplotypes. For recombinant genes for which short-read sequencing does not enable to reconstruct the haplotype, such as the *cid* genes used here, Nanopore sequencing of PCR products enables to have access to the haplotype in a single read.

We highlight the numerous chimeric variants created by PCR. Our results confirm previous results showing that the probability of chimera formation increases with the number of cycles ([28], here with 31 vs 35 cycles), and when similar template sequences are amplified in the same PCR reaction [28,31,32]. Recently, along with the increased sequencing of PCR products with the rise of metabarcoding, and of other methods such as MPRAs (massively parallel reporter assays), some studies examined the impact of such chimeras on multiplexing results, showing that up to 28% of the sequences correspond to mistags, resulting from tag-switching events [33]. Finding such high occurences of PCR-linked errors made previous studies seek for the optimal PCR conditions to reduce chimera formation, examining for instance the consequences of the type of polymerase used [34] or various factors such as the number of cycles used or the amount of DNA template [35,36]). Here, we further stress the urge to consider such chimeras, especially when working on recombinant genes, since the genetic architecture of those genes makes harder to discriminate true variants and chimeras.

PCR-induced recombinations being an issue regardless of the sequencing method, Nanopore sequencing of multigene families has massive advantages compared to cloning and Sanger sequencing of PCR products: it enables to sequence longer fragments (up to 3kb here) which could hardly be inserted into plasmids, and it is much more time effective as multiple genes can be sequenced in the same sequencing well. While we sequenced just two genes at the same time here, simultaneous acquisition of multiple PCR products through Nanopore sequencing has already been validated, increasing the cost-effectiveness of the method.

Since we found that Nanopore sequencing coverage strongly differed among variants, we sought to see if these coverage differences could be reproduced using other methods, or if they likely resulted from Nanopore sequencing biases – those biases being numerous, see [4] for a review. We confirmed that coverage variations in Nanopore sequencing could give hints of copy number variations using digital droplet PCR (ddPCR). To go from coverage variation to variant copy number, the total number of *cid* copies is required. This can be easily obtained with a generic quantitative PCR normalized on a single-copy gene. While ddPCR mirrors Nanopore sequencing coverage, it has to be kept in mind that both methods involve a PCR, and that PCRs have been shown to skew template-to-product ratios, with a bias towards some gene versions [37]. Some gene variants may be more easily amplifiable, resulting in higher coverage. In making controls to ensure that all gene copies were amplified in ddPCR, we found that fragment size, and by extent reaction efficacy, unexpectedly influenced the outcomes of ddPCR, which could result in some biases if one was to compare concentrations of different genes using amplicons differing in size. This bias can be counterbalanced by ensuring that all amplified fragments have approximately the same size, but has to be carefully taken into account when using ddPCR.

While the method presented here requires a good pre-existing reference database, the recent development of methods to work on multigene families based on less error-prone PacBio sequencing data [5], along with the improvement of both Nanopore basecalling [7,38] and error-correction [9] suggest that *de novo* assembly of multigene families will be possible with Nanopore sequencing in the coming years. Overall, in spite of some limitations due to the PCR step itself, this method enables a quick acquisition of numerous variants, which could be further increased by acquiring more than two genes at the same time. This paves the way to wide scale multigene family acquisitions and, with the present study, to a better understanding of links between *cid* genes and cytoplasmic incompatibility phenotypes.

## 4. Methods

For all experiments, total DNA was extracted on adult mosquitoes following the acetyltrimethilammonium bromide (CTAB) protocol [39].

### 4.1. Mosquito lines used in this study

All the mosquito lines used were isofemale lines, i.e. lines obtained by rearing the progeny of a single female. *cidA* and *cidB* repertoires were acquired through Nanopore sequencing of PCR products for 13 isofemale lines reared at the laboratory (Table S5).

All isofemale lines were reared in 65 dm^3^ screened cages, in a single room maintained at 26°C, under a 12h light/ 12h dark cycle. Larvae were fed with a mixture of shrimp powder and rabbit pellets, and adults were fed on honey solution. Females were fed with turkey blood, using a Hemotek membrane feeding system (Discovery Workshops, UK), to enable them to lay eggs.

### 4.2. Cloning and Sanger sequencing

Amplifications of the variable parts of the *cidA* and *cidB* genes were done using the primer pairs 1/2 and 7/8 [17], giving respectively 1,300 bp and 1,264 bp PCR products (Table S3). Amplifications were done using the GoTaq polymerase (Promega).

PCR products were cloned following the procedure from [17].

### 4.3. Sequence acquisition through Nanopore sequencing

*cid* genes were amplified using the same primer pairs as for Sanger sequencing, on DNA extracted from a single adult mosquito. For each of the strains, repertoires were acquired for two distinct individuals. PCR products were purified using CleanPCR beads at 1.8X (CleanNA) and quantified using a Qubit fluorometer and Qubit DS DNA Broad Range kits (ThermoFisher). Purified PCR products of each gene were pooled in an equimolar mix. The first runs were sequenced using a Nanopore protocol that involved a further amplification step. Since we found that amplifications create artificial recombinations, the following sequences were obtained using a PCR-free protocol, by the MGX platform (Montpellier GenomiX). The DNA amplicons were quantified using a Tecan infinite 500 Fluorometer (Tecan, Switzerland) with a dsDNA High Sensitivity Assay Kit (Thermo Fisher Scientific, Massachusetts, USA). The fragments size distribution was checked using a 5200 Fragment analyzer (Agilent, USA) system with a Sandard NGS kit (Agilent, USA). The amplicons libraries construction was done according the Nanopore protocole NBA_9102_V109_revA_09Jul2020. Two hundred nanograms of DNA are end repaired and dA-tailed using NEBNext End repair/dA-tailing Module (E7546, New England Biolabs, Ipswich, Massachusetts, USA). The samples were then barcoded using EXP-NBD196 (barcodes 1–96) kits (Oxford Nanopore Technologies, Oxford, UK) and ligation Sequencing Kit 1D SQK-LSK109, (Oxford Nanopore Technologies, Oxford, UK). Barcoded samples were pooled and purified using 0.4 volume of AMPure XP magnetic beads. The AMII sequencing adapter (Oxford Nanopore Technologies, Oxford, UK) was ligated to barcoded pools using Quick T4 DNA ligase (New England Biolabs, Ipswich, Massachusetts, USA) and the sequencing libraries were purified using 0.4 volume of AMPure XP magnetic beads. MinION sequencing was performed as per manufacturer’s guidelines using R9.4.1 flow cells FLO-MIN106, ONT and controlled using Oxford Nanopore Technologies MinKNOW software version v20.06.5. Flow cells were then transferred to Nanopore MinION Mk1b (Oxford Nanopore Technologies, UK) for Nanopore single molecular sequencing.

Base calling was performed after sequencing using the GPU-enabled guppy basecaller in high accuracy mode for 96 samples (version 4.2.2), 96 samples (version 4.4.2), 74 samples (version 5.0.14) and 96 samples (version 5.0.16). A quality control was done using PycoQC [40] (version v2.5.2) for all samples.

### 4.4. Establishment of an artificial reference base

The *cid* genes are composed of two variable regions separated by a monomorphic region, and recombination occurs between those two regions (Fig 1, [17]). We thus completed the pool of references already sequenced by creating an *in silico* pool of references for each of the five wPip group [41] by combining all the previously sequenced upstream and downstream regions within each group.

### 4.5. Pipeline details

Throughout the whole pipeline, mapping was done using minimap2 [21], with parameters - ax map-ont, unless specified otherwise. We first separated *cidA* and *cidB* reads by mapping all reads on a short monomorphic sequence of each gene, corresponding to positions 228 to 327 and 1211 to 1309 for *cidA* and *cidB* respectively. After this step, there were around 15,000-20,000 reads per gene. The pipeline was then run separately for *cidA* and *cidB* reads, the following steps of the pipeline being identical for both genes.

For each gene, reads were mapped on the full pool of references and secondary alignments were removed. Samtools coverage was then used (samtools 1.15, [22]) to obtain the coverage of each reference (Fig 1). Using specific PCRs, we determined that selecting references which had a coverage above 10% of the total number of reads enabled to get all the true references, and excluded artifactual chimeric reads (see below). References above that threshold were extracted, along with reads mapping on these references. Since highly similar references artificially introduce INDELs, and decrease coverage in SNP calling when using bcftools mpileup, we looked for pairs of references differing by less than 3 SNPs and kept a single representative for each pair (Fig 1). Exclusion was done by computing the raw distance among sequences using the dist.dna function in the R package ape [42].

Reads which had mapped on at least a reference were then mapped on the reference subset, and SNPs were called using bcftools mpileup (with the ont config, a minimum mapping quality of 10, a disabled BAQ and a max depth of 75,000) and bcftools call (multiallelic caller, with an expected substitution rate of 0.5). If no SNPs were called, variants present and their respective coverage were extracted to assemble the repertoire. If SNPs were called, two alternative cases occurred: (i) cases where haplotypes could be directly deduced from the SNP calling or (ii) cases where haplotype calling was required to sort out haplotypes. In the first case, the consensus sequence was obtained using bcftools consensus (option --haplotype I), while in the second, WhatsHap phase [23] was used to phase the haplotypes. If the respective coverage of the references and alternative gene version can vary, we never found more than two distinct alleles at a given position, and thus used the default WhatsHap settings (in cases where more than 2 distinct variants can correspond to a given known reference, WhatsHap polyphase could be used).

### 4.6. Nanopore read simulation

Our read simulation script is based on the simulation of full-length transcripts used in [9] with modifications to fit our targeted sequencing scenario. We simulate reads as follows. The script takes as input a fasta file with a number of *N* starting reference sequences, an integer *C* (*C>N*) of targeted simulated references, a fraction S corresponding the mutation rate of the references, and the mean error rate X of the reads. Furthermore, the script has two settings regarding mutation type. It can simulate substitutions only, or both SNPs and indels. The outputs of the script are a fasta file with the references (original and simulated) from which the reads were simulated, and a fastq file with the simulated reads. We now describe the workflow of the script.

First, *C* references are simulated from the *N* starting references as follows. The *N* original references are added to a pool of simulated references here denoted *R*. Then, a random reference *r* is sampled from *R* and mutated with the mean mutation rate *S* according to the mutation profile specified and placed back into the pool *R* which now contains *N*+1 references. This procedure is repeated until the pool *R* contains *C* references. Note that a simulated reference can be selected from *R* and in turn be mutated into a new reference, creating a tree-like evolution structure.

Second, reads are sampled from the pool of *C* references in the same way as in [9] for full length transcripts. We briefly describe the procedure here, for details see Supplementary note 1 in [9]. To simulate a read we pick a reference in *R* at random. We simulate a quality value uniformly at random over each base pair in the sampled reference. The base is assigned the Phred score and we introduce an error at that position with a probability corresponding to the phred score. The error types are either deletion, substitution, or insertion with probabilities of 0.45, 0.35, and 0.2, respectively, which roughly mimics the error profile of ONT data although nanopore base calling algorithms changes rapidly. Our script is able to produce reads with a mean error of ~3.9%, 7, and ~11.4% error rate through different ratios of phred quality values.

### 4.7. Testing the coverage of specific variants using ddPCR

Prior to the experiment, *cidA* variants of the Lavar strains were cloned as described above and in [17]. Clones corresponding to each variant were used as controls for the PCR specificity, using pairs of specific primers that should amplify specifically a single clone. Primer pairs used are shown in Table S4.

The digital PCR assays were then performed using the Naica digital PCR system (Stilla Technologies). The dPCR reaction mixture (25 μL) contained 5 μL of Quantabio PerfeCTa Multiplex qPCR ToughMix 5x (Quantabio), 1 μL of Dextran Alexa Fluor 647 10,000 MW (Thermofisher), 1.9 μL of Evagreen 20x (Biotium), 3.125 μL of the target primer set (final concentration of 125nM) and nuclease-free water up to 25 μL. DNA was added in a quantity sufficient to get enough positive droplets (for full mosquitoes, DNA amount could not be quantified as infection levels of the endosymbiotic *Wolbachia* vary and extracted DNA is always mixed with host DNA). The reaction mixtures were loaded into wells of Sapphire chip and were subsequently emulsified (20,000 to 30,000 droplets/sample) and amplified in a geode thermocycler (Stilla technologies). The ddPCR conditions used were 10 min initial denaturation at 95°C, followed by 45 cycles of 95°C for 30s, 58°C for 15s and 72°C for 30s. After template amplification, the chips were transferred to the reader. Extracted fluorescence values for each droplet were analyzed using the Crystal Miner software (Stilla Technologies).

## Code availability

All scripts used for simulating datasets, to run the pipeline and its evaluation are found at https://github.com/alnam3/nanoseq. Specifically, the source code of the pipeline are found under https://github.com/alnam3/nanoseq/Mainpipeline, the code to simulate Nanopore reads under https://github.com/alnam3/nanoseq/Nano-read-simulator, and the suggested awk script to test for missed SNPs under https://github.com/alnam3/nanoseq/MissedSNPs.

## Acknowledgements

We thank the MBB platform for their help on code-related issues. Di91al PCRs were run on the ISEM qPCR platform. Qubit were run using GenSeq platform facilities. This project was funded by the MUSE project with the reference ANR-16-IDEX-0006. Kristoffer Sahlin was supported by the Swedish Research Council (SRC, Vetenskapsrådet) under Grant No. 2021-04000.

We thank Nicole Pasteur for thorough reading and helpful comments on previous versions of this manuscript.

## References

1. Magadum S, Banerjee U, Murugan P, Gangapur D, Ravikesavan R. 2013 Gene duplication as a major force in evolution. J. Genet. 92, 155–161. (doi:10.1007/S12041-013-0212-8)

2. Gresham D, Dunham MJ, Botstein D. 2008 Comparing whole genomes using DNA microarrays. Nat. Rev. Genet. 9, 291–302. (doi:10.1038/nrg2335)

3. Assogba BS et al. 2016 The *ace-1* Locus Is Amplified in All Resistant Anopheles *gambiae* Mosquitoes: Fitness Consequences of Homogeneous and Heterogeneous Duplications. PLoS Biol. 14, 1–26. (doi:10.1371/journal.pbio.2000618)

4. Delahaye C, Nicolas J. 2021 Sequencing DNA with nanopores: Troubles and biases. PLoS One 16. (doi:10.1371/journal.pone.0257521)

5. Sahlin K, Tomaszkiewicz M, Makova KD, Medvedev P. 2018 Deciphering highly similar multigene family transcripts from Iso-Seq data with IsoCon. Nat. Commun. 9, 1–12. (doi:10.1038/s41467-018-06910-x)

6. Loman NJ, Quick J, Simpson JT. 2015 A complete bacterial genome assembled de novo using only nanopore sequencing data. Nat. Methods 2015 128 12, 733–735. (doi:10.1038/nmeth.3444)

7. Ahsan MU, Liu Q, Fang L, Wang K. 2021 NanoCaller for accurate detection of SNPs and indels in difficult-to-map regions from long-read sequencing by haplotype-aware deep neural networks. Genome Biol. 22, 1–33. (doi:10.1186/s13059-021-02472-2)

8. Nowak A, Murik O, Mann T, Zeevi DA, Altarescu G. 2021 Detection of single nucleotide and copy number variants in the Fabry disease-associated GLA gene using nanopore sequencing. Sci. Rep. 11, 1–7. (doi:10.1038/s41598-021-01749-7)

9. Sahlin K, Medvedev P. 2021 Error correction enables use of Oxford Nanopore technology for reference-free transcriptome analysis. Nat. Commun. 12, 1–13. (doi:10.1038/s41467-020-20340-8)

10. Whitford W, Hawkins V, Moodley KS, Grant MJ, Lehnert K, Snell RG, Jacobsen JC. 2022 Proof of concept for multiplex amplicon sequencing for mutation identification using the MinION nanopore sequencer. Sci. Rep. 12, 1–9. (doi:10.1038/s41598-022-12613-7)

11. Beckmann JF, Ronau JA, Hochstrasser M. 2017 A *Wolbachia* deubiquitylating enzyme induces cytoplasmic incompatibility. Nat. Microbiol. 2. (doi:10.1038/nmicrobiol.2017.7)

12. LePage DP et al. 2017 Prophage WO genes recapitulate and enhance *Wolbachia*-induced cytoplasmic incompatibility. Nature 543, 243–247. (doi:10.1038/nature21391)

13. Zabalou S, Riegler M, Theodorakopoulou M, Stauffer C, Savakis C, Bourtzis K. 2004 Wolbachia-induced cytoplasmic incompatibility as a means for insect pest population control. Proc. Natl. Acad. Sci. U. S. A. 101, 15042–15045. (doi:10.1073/pnas.0403853101)

14. Atyame CM et al. 2011 Cytoplasmic incompatibility as a means of controlling culex pipiens quinquefasciatus mosquito in the islands of the South-Western Indian Ocean. PLoS Negl. Trop. Dis. 5, 20–22. (doi:10.1371/journal.pntd.0001440)

15. Nikolouli K et al. 2018 Sterile insect technique and Wolbachia symbiosis as potential tools for the control of the invasive species Drosophila suzukii.J. Pest Sci. (2004). 91, 489–503. (doi:10.1007/s10340-017-0944-y)

16. Utarini A et al. 2021 Efficacy of Wolbachia-Infected Mosquito Deployments for the Control of Dengue. N. Engl. J. Med. 384, 2177–2186. (doi:10.1056/nejmoa2030243)

17. Bonneau M, Atyame CM, Beji M, Justy F, Cohen-Gonsaud M, Sicard M, Weill M. 2018 *Culex pipiens* crossing type diversity is governed by an amplified and polymorphic operon of *Wolbachia*. Nat. Commun. 9, 319. (doi:10.1038/s41467-017-02749-w)

18. Bonneau M, Caputo B, Ligier A, Caparros R, Unal S, Perriat-Sanguinet M, Arnoldi D, Sicard M, Weill M. 2019 Variation in *Wolbachia cidB* gene, but not *cidA*, is associated with cytoplasmic incompatibility *mod* phenotype diversity in *Culex pipiens*. Mol. Ecol. 28, 4725–4736. (doi:10.1111/mec.15252)

19. Bonneau M, Landmann F, Labbé P, Justy F, Weill M, Sicard M. 2018 The cellular phenotype of cytoplasmic incompatibility in Culex pipiens in the light *of cidB* diversity. PLoS Pathog. 14, e1007364. (doi:10.1371/journal.ppat.1007364)

20. Sicard M, Namias A, Perriat-Sanguinet M, Carron E, Unal S, Altinli M, Landmann F, Weill M. 2021 Cytoplasmic incompatibility variations in relation with *Wolbachia cid* genes divergence in *Culex pipiens*. MBio 12, e02797–20. (doi:10.1128/mBio.02797-20)

21. Li H. 2018 Minimap2: Pairwise alignment for nucleotide sequences. Bioinformatics 34, 3094–3100. (doi:10.1093/bioinformatics/bty191)

22. Danecek P et al. 2021 Twelve years of SAMtools and BCFtools. Gigascience 10, 1–4. (doi:10.1093/gigascience/giab008)

23. Martin M, Patterson M, Garg S, Fischer SO, Pisanti N, Gunnar W, Marschall T. 2016 WhatsHap□: fast and accurate read-based phasing., 1–18.

24. Yang C, Chu J, Warren RL, Birol I. 2017 NanoSim: Nanopore sequence read simulator based on statistical characterization. Gigascience 6, 1–6. (doi:10.1093/gigascience/gix010)

25. Li Y, Han R, Bi C, Li M, Wang S, Gao X. 2018 DeepSimulator: A deep simulator for Nanopore sequencing. Bioinformatics 34, 2899–2908. (doi:10.1093/bioinformatics/bty223)

26. Stöcker BK, Köster J, Rahmann S. 2016 SimLoRD: Simulation of Long Read Data. Bioinformatics 32, 2704–2706. (doi:10.1093/bioinformatics/btw286)

27. Faucon PC, Balachandran P, Crook S. 2017 SNaReSim: Synthetic Nanopore Read Simulator. Proc. - 2017 IEEE Int. Conf. Healthc. Informatics, ICHI 2017, 338–344. (doi:10.1109/ICHI.2017.98)

28. Smyth RP, Schlub TE, Grimm A, Venturi V, Chopra A, Mallal S, Davenport MP, Mak J. 2010 Reducing chimera formation during PCR amplification to ensure accurate genotyping. Gene 469, 45–51. (doi:10.1016/j.gene.2010.08.009)

29. Di Giallonardo F et al. 2013 Next-Generation Sequencing of HIV-1 RNA Genomes: Determination of Error Rates and Minimizing Artificial Recombination. PLoS One 8, e74249. (doi:10.1371/JOURNAL.PONE.0074249)

30. Liu J, Song H, Liu D, Zuo T, Lu F, Zhuang H, Gao F. 2014 Extensive Recombination Due to Heteroduplexes Generates Large Amounts of Artificial Gene Fragments during PCR. PLoS One 9, e106658. (doi:10.1371/JOURNAL.PONE.0106658)

31. Judo MSB, Wedel AB, Wilson C. 1998 Stimulation and suppression of PCR-mediated recombination. Nucleic Acids Res. 26, 1819. (doi:10.1093/NAR/26.7.1819)

32. Fonseca VG, Nichols B, Lallias D, Quince C, Carvalho GR, Power DM, Creer S. 2012 Sample richness and genetic diversity as drivers of chimera formation in nSSU metagenetic analyses. Nucleic Acids Res. 40, e66–e66. (doi:10.1093/NAR/GKS002)

33. Esling P, Lejzerowicz F, Pawlowski J. 2015 Accurate multiplexing and filtering for high-throughput amplicon-sequencing. Nucleic Acids Res. 43, 2513–2524. (doi:10.1093/nar/gkv107)

34. Nagai S, Sildever S, Nishi N, Tazawa S, Basti L, Kobayashi T, Ishino Y. 2022 Comparing PCR-generated artifacts of different polymerases for improved accuracy of DNA metabarcoding. Metabarcoding Metagenomics 6 e77704 6, e77704-. (doi:10.3897/MBMG.6.77704)

35. Potapov V, Ong JL. 2017 Examining sources of error in PCR by single-molecule sequencing. PLoS One 12, 1–19. (doi:10.1371/journal.pone.0169774)

36. Omelina ES, Ivankin A V., Letiagina AE, Pindyurin A V. 2019 Optimized PCR conditions minimizing the formation of chimeric DNA molecules from MPRA plasmid libraries. BMC Genomics 20, 1–10. (doi:10.1186/S12864-019-5847-2/TABLES/3)

37. Polz MF, Cavanaugh CM. 1998 Bias in Template-to-Product Ratios in Multitemplate PCR. Appl. Environ. Microbiol. 64, 3724. (doi:10.1128/AEM.64.10.3724-3730.1998)

38. Rang FJ, Kloosterman WP, de Ridder J. 2018 From squiggle to basepair: Computational approaches for improving nanopore sequencing read accuracy. Genome Biol. 19, 1–11. (doi:10.1186/s13059-018-1462-9)

39. Rogers SO, Bendich AJ. 1994 Extraction of total cellular DNA from plants, algae and fungi. In Plant Molecular Biology Manual, pp. 183–190. Springer Netherlands. (doi:10.1007/978-94-011-0511-8_12)

40. Leger A, Leonardi T. 2019 pycoQC, interactive quality control for Oxford Nanopore Sequencing. J. Open Source Softw. 4, 1236. (doi:10.21105/JOSS.01236)

41. Atyame CM, Delsuc F, Pasteur N, Weill M, Duron O. 2011 Diversification of *Wolbachia* endosymbiont in the *Culex pipiens* mosquito. Mol. Biol. Evol. 28, 2761–2772. (doi:10.1093/molbev/msr083)

42. Paradis E et al. 2022 Package ‘ape’ Title Analyses of Phylogenetics and Evolution Depends R (>= 3.2.0). R Top. Doc.

