## Supplementary figures and images for "Nanopore sequencing enables multigenic family reconstruction despite highly frequent PCR-induced recombination"

### Supplementary Figure 1

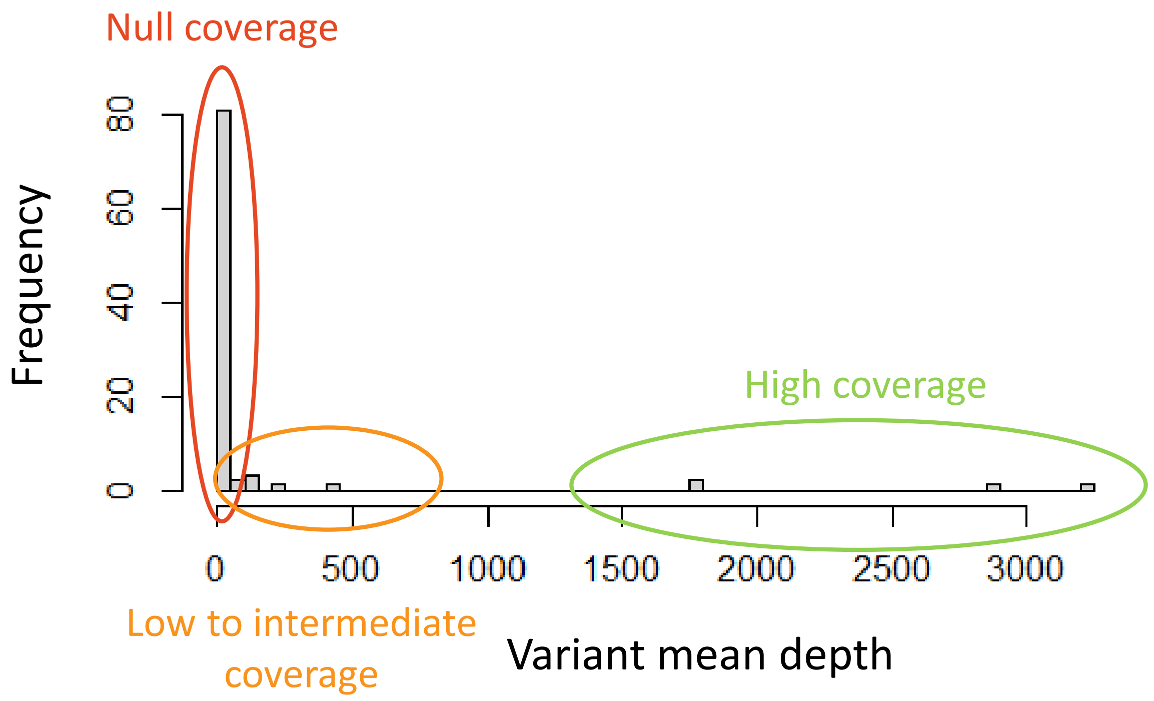
