## Supplementary Legends for "Nanopore sequencing enables multigenic family reconstruction despite highly frequent PCR-induced recombination"

Figure S1. Example of the coverage depth distribution obtained on a single individual, when mapping the reads on all the possible references. Few variants are well covered and are undoubtedly present (circled in green), most variants have a null coverage and are absent (circled in red), and some variants have a low to intermediate coverage (circled in orange). It is to determine whether these last variants were true or artifactual that specific PCRs were used (Fig. 2).

Table S1. *cidA* and *cidB* repertoires of the strains sequenced by both Sanger and Nanopore sequencing methods. Colors and text show (dis)agreement between the two sequencing methods, along with results on presence/absence of variants obtained through specific targeted PCRs.

Table S2. Testing digital droplet PCR (ddPCR) quantifications of *cidA* variants from the Lavar *w*Pip genome in controlled conditions. As specific primers for each *cidA* variants present in the Lv strain are ~600 bp, qPCR could not be used to verify whether variations in coverage observed with Nanopore sequencing were relevant for estimating copy numbers.

**A**. ddPCR quantifications of each individual variant using specific primers on a same DNA quantity from each clone. Note the similar results are similar, indicating that there was no efficiency difference in PCR amplification.

**B**. ddPCR quantification of an equimolar mix of DNA from the three cloned variants using generic *cidA* primer pairs that amplify 150, 580 and 1300 bp fragments. Surprisingly, the size of the generic fragment strongly influences the obtained copy numbers. For further experiments, we used the generic *cidA* primers giving fragments the size of which was the closest to the specific fragments, since it gave the quantification closest to the expected copy number (586.6 being the sum of all specific variant concentrations).

Concentrations are given in copies/μL, along with a 95% confidence interval.

Table S3. Comparison of the respective coverage of *cidA* variants of the Lavar strain obtained with Nanopore sequencing of *cidA* PCR products and with digital droplet PCR (ddPCR) on total DNA from Lavar wPip strain. For ddPCR, concentrations are given in copies/μL along with a 95% confidence interval. ddPCR was also run with a generic *cidA* to ensure that no *cidA* copy was missed. Relative coverage of the variants are similar in Nanopore sequencing and in ddPCR.

Table S4. Primer pairs used for digital droplet PCR.

Table S5. Origin and reference of all strains used in this study.
